# Species-specific dynamics may cause deviations from general biogeographical predictions – evidence from a population genomics study of a New Guinean endemic passerine bird family (Melampittidae)

**DOI:** 10.1101/2023.10.19.563072

**Authors:** Ingo A. Müller, Filip Thörn, Samyuktha Rajan, Per Ericson, John P. Dumbacher, Gibson Maiah, Mozes Blom, Knud A. Jønsson, Martin Irestedt

## Abstract

New Guinea, the largest tropical island, is topographically complex and is dominated by a large central mountain range surrounded by multiple smaller isolated mountain regions along its perimeter. The island is biologically hyper-diverse and harbours an avifauna with many species found only there. The family Melampittidae is endemic to New Guinea and consists of two monotypic genera: *Melampitta lugubris* (Lesser Melampitta) and *Megalampitta gigantea* (Greater Melampitta). Both Melampitta species have scattered and disconnected distributions across New Guinea in the central mountain range and in some of the outlying ranges. While *M. lugubris* is common and found in most montane regions of the island, *M. gigantaea* is elusive and known from only six localities in isolated pockets on New Guinea with very specific habitats of limestone and sinkholes. In this project, we apply museomics to determine the population structure and demographic history of these two species. We re-sequenced the genomes of all seven known *M. gigantaea* samples housed in museum collections as well as 24 *M. lugubris* samples from across its distribution. By comparing population structure between the two species, we investigate to what extent habitat dependence, such as in *M. gigantaea*, may affect population connectivity. Phylogenetic and population genomic analyses, as well as acoustic differentiation, revealed that *M. gigantaea* consists of a single population in contrast to *M. lugubris* that shows much stronger population structure across the island. This work sheds new light on the mechanisms that have shaped the intriguing distribution of the two species within this family and is a prime example of the importance of museum collections for genomic studies of poorly known and rare species.

## Introduction

What determines the build-up of biodiversity in space and through time is a long-standing question within biology. The accumulation of phenotypic and genetic differences between populations can only be generated through reproductive isolation that impedes genetic exchange between populations [1,2]. More explicitly, gene flow between diverging populations must be sufficiently limited so that genetic exchange does not exceed the accumulation of differentiation. Barriers underlying reproductive isolation, may differ markedly. They may be postzygotic and arise from genetic incompatibilities, which produce hybrid offspring that have either reduced fitness or are infertile [3–5]. Alternatively, barriers may be prezygotic and decrease the probability of mating events between populations, due to mating preferences, or through geographical (allopatric) or habitat barriers that separate different populations [4–6].

Mountains represent a classic example of geographical barriers both as physical barriers for populations but also because they harbour highly differentiated environments at different elevations. For sedentary lowland populations, mountains may represent unsurpassable barriers, which may over time lead to isolation and differentiation of separate lowland populations. Evidence for such montane barriers restricting gene flow between lowland populations are known from various organismal groups such as amphibians, spiders and coniferous trees [7–9]. Alternatively, extensive lowland valleys can also act as barriers to geneflow between populations adapted to high elevations. Lowland environments may be unsuitable for such mountain-adapted individuals, which over time become isolated on a series of mountaintops or “sky islands” [10,11] as known from some groups of birds, lizards and plants [12–16].

Related to this is the observation that older taxa are often found at higher elevations, while young lineages that are generally widespread, good dispersers and show little differentiation inhabit the lowlands (e.g. 16–19). Such observations (mostly from island systems) have led to the formulation of the concept of taxon cycles, in which taxa pass through phases of expansions and contractions. The concept predicts that over time, taxa move into high elevation habitats either because they are outcompeted by new young taxa in their original (lowland) habitats or because they specialise and adapt to new environments at higher elevations [17,18,20–22].

Recent work on the New Guinean avifauna has provided empirical evidence in favour of species originating in the lowlands from where they move into the highlands over time and become relictual specialists [16,22,23], although some colonisation from mountaintop to mountaintop has also been shown to occur [15]. In addition, recent Pleistocene speciation events on New Guinea are mainly the result of changes in habitat distributions due to climate fluctuations, as this has caused species with continuous distributions to become geographically fragmented [24–26]. Pliocene speciation events, on the other hand, are driven mainly by geological processes such as montane uplift, which is known to have caused barriers to gene flow [27–30].

The Melampittidae represents an example of an old passerine family with only two deeply diverged species in monotypic genera. Their taxonomic affinities have been difficult to establish, but recent genetic results have placed the family as sister to crows (Corvidae) and shrikes (Laniidae) with an estimated divergence time from these at ca. 17 Mya [31]. One of the species, *Melampitta lugubris* (Lesser Melampitta) is relatively common at high elevations (1150-3500 m asl.), in accordance with the notion that older species tend to occupy higher elevations [21,22,32]. The other species, *Megalampitta gigantea* (Greater Melampitta) is only known from six localities at mid-elevations (650 −1400 m asl.) scattered across New Guinea. Based on few field observations, it is considered to be sedentary and to have limited flight capabilities [33,34]. Within its range, *M. gigantea* is associated with very specific karstic habitats where it has been observed to spend considerable time nesting in narrow limestone sinkholes in which the birds have to climb in and out [33]. In contrast to *M. lugubris* the distribution of *M. gigantea* does not fit the general pattern that old taxa tend to occupy higher elevation.

In this genomic study we determine the population structure within *M. lugubris* and *M. gigantea* to understand how habitat connectivity across space and through time has shaped differentiation in these two species. Based on the contemporary distributions of the two species we hypothesize that:

1. The 7 individuals of *M. gigantea* represent several distinct evolutionary entities/populations, as the species is a poor disperser and has a fragmented distribution across New Guinea where it is associated with specific karst limestone habitat with sinkholes.
2. Individuals of *M. lugubris* represent a relatively cohesive group, yet with some population structure as deep lowland valleys may prevent gene flow between the various montane populations in the Central Range, the Huon mountains in the northeast and the Arfak mountains in the northwest.

## Material & Methods

### DNA sampling, sequencing and read processing

In this study, we follow the taxonomy of the IOC World Bird List [35]. We sampled 24 individuals of *Melampitta lugubris* of which 22 were footpads from museum specimens and two were fresh tissue samples. Additionally, we sampled 7 individuals of *Megalampitta gigantaea*, which represent all known samples present in museum collections. One of these samples was a fresh blood sample. The rest were footpads from historical samples (for a detailed list of samples and the museum collections in which they are stored see S1 Table). The work is mainly based on old museum specimens for which the Nagoya Protocol does not apply. The few fresh tissue samples included in the study are from already preserved samples at natural history museums, for which all required permits are available.

DNA from fresh blood/tissue samples was extracted using Qiagen’s DNeasy Blood and Tissue kits. For DNA extraction and sequencing library preparation of historical samples, we followed a modified version of Meyer and Kircher [36] that proved suitable for avian museum samples [37]. In short, we extract DNA from toepad tissue mainly following the instructions from Qiagen for animal tissue with the addition of Dithiothreitol (DTT) to improve the ligation yield. During library preparation, we treat our samples with USER enzyme to reduce deamination patterns that are typical for fragmented DNA from historical or ancient samples [38]. For a detailed protocol see [37]. Whole genome re-sequencing was performed on Illumina NovaSeq 6000 machines on S4 flow cells going through 200 cycles with a read length of 2 x 100 bp at the National Genomics Infrastructure (NGI) in Stockholm. Up to 24 samples with four libraries each were multiplexed on a single flow cell lane.

Sequenced reads were then polished using the reproducible *Nextflow* workflow *nf-polish (*https://github.com/MozesBlom/nf-polish) [39,40] (see S2 Table for specific github commits used). This pipeline performs multiple polishing steps, including deduplication, adapter- and quality-based trimming, read merging and the removal of low-complexity reads. Polished reads were then mapped onto a reference genome using *nf-μmap* (https://github.com/IngoMue/nf-umap) applying bwa-mem2 as mapping algorithm and which also allowed us to evaluate the quality control after mapping and investigate damage patterns that are typical for historical DNA [41,42]. We used the hooded crow (*Corvus cornix*, Refseq GCF_000738735.5, [43]) as our reference genome as it represents a reasonable closely related species with a high-quality chromosome level assembly [31].

### Phylogenetic analyses on mitochondrial and nuclear DNA

To assemble the full mitogenome from our polished reads we used nf_mito-mania with default settings (https://github.com/FilipThorn/nf_mito-mania) [44]. Variant calling implemented in this pipeline filters sites with a depth-of-coverage below 20 or above three times the average depth-of-coverage across the whole mitogenome of each individual. The resulting consensus sequences of every individual were aligned using *MAFFT* (v7.407, [45], see S2 File for specific flags). An occasional artefact of Mitobim where mitochondrial assemblies become longer than they are supposed to be resulted in overhanging sequences in some individuals. These overhangs were then cut out of the alignment after visual inspection using *Geneious Prime 2023.0.4* so that the final alignment consisted only of overlapping reads (total length 17 112 bp including gaps), which were then used as input for *RAxML-NG* (v1.1.0, [46] using the GTR+G substitution model, 100 bootstrapping replicates and ten randomized parsimony starting trees to generate a mitochondrial phylogenetic tree. Mitochondrial assemblies were also forced into diploid variant calls to check for contamination in our samples. As mitochondria are haploid, heterozygote sites are not expected and could therefore be indicative of cross-contamination.

For the nuclear phylogenetic tree, we used the previously mapped .bam files excluding individuals with very low mean depth-of-coverage (n = 3, DoC < 4 *x*) to call variants for each individual using *freebayes* (v1.3.1-dirty, [47]). Polymorphic sites were filtered based on their quality score (> 20), allelic balance (≥ 0.2), and minimum and maximum depth-of-coverage (3 *x* / 100 *x*). We also decomposed multiple nucleotide polymorphisms (MNPs) into single nucleotide polymorphisms (SNPs) and masked heterozygous positions and indels. These filtered .vcf files were then used as input files for the reproducible *Nextflow* workflow *nf-phylo* with default settings (https://github.com/MozesBlom/nf-phylo) [48] to generate both concatenated (*IQtree2*) and summary coalescent (*ASTRAL3*) species trees based on ‘gene’ trees of different window-sizes (2000, 5000, 10000, 20000 base pairs).

### Population structure and differentiation

To quantify population substructure and to estimate levels of differentiation between samples, we used a genotype likelihood approach as implemented in ANGSD (v0.938) as this is better suited for low coverage data [49]. Specific commands and the filters used are explained in the S2 File. The filters we used for admixture and principal component analyses (PCAs) were slightly different from those used to calculate nucleotide diversity, heterozygosity and Tajima’s D. As we did not have an ancestral genome available, we used the reference genome as ancestral sequence and folded the site frequency spectra (SFS). PCAs were performed through *PCAngsd* [50] and plotted with custom R scripts through RStudio (v 2023.03.0 build 386, R version 4.1.1, [51,52]). Admixture analyses to determine population structure was run through *NgsAdmix* [53] running up to K = 10 with ten replicates for each K and visualised with custom R scripts. Individual heterozygosity was estimated by generating a site frequency spectrum for each individual and dividing the number of sites with one derived allele divided by the total number of sites as performed by e.g. Hansen et al. [54]. Using SFS for each species, nucleotide diversity and Tajima’s D were both estimated for each chromosome as well as in 20 kb windows sliding in steps of 10 kb using the *thetaStat* command. We divided the pairwise theta estimator (*tP*) by the total number of Sites (*nSites*) of each chromosome/window to calculate nucleotide diversity. Statistical significance of differences in heterozygosity and nucleotide diversity between the two species was checked using Welch’s t-test after verifying normal distributions and inequal variances within the data.

### Estimation of effective population sizes through time and divergence times

To estimate effective population sizes through time, we used pairwise sequentially Markovian coalescent (PSMC) [55] (for details on the method see S1 file). As an estimate of the neutral genomic mutation rate per generation we used 4.6*10^-9^ as obtained in a study of the collared flycatcher *Ficedula albicollis* [56]. We set the estimated generation time for *M. lugubris* to 3.90 years and for *M. gigantea* to 4.58 years [57]. The parameters for the PSMC analysis were set to “ -N30 -t5 -r5 -p 4 + 30*2 + 4 + 6 + 10” following Nadachowska-Brzyska et al. [58]. The authors observed no significant change in curve shape when modifying the atomic vectors parameter (-p) and applied the same settings to several different avian species. We only ran PSMCs for the two samples of *M. gigantea* with the highest depth-of-coverage and for each of the five identified clusters within *M. lugubris* (West, Central, East, Huon, Southeast). False negative rates (FNRs) were adjusted based on depth-of-coverage. If depth-of-coverage was higher than 15 X, FNR was kept at 0. However, if individual A had a depth-of-coverage higher than 15 X and individual B had a depth-of-coverage below 15 X, then individual B would have an FNR of 0.1 + 0.1 * x, where x is the depth-of-coverage of individual A divided by depth-of-coverage of individual B. If both individuals had a depth-of-coverage < 15 X, we used 0.1 for both individuals.

To estimate divergence times between the two species, but also between the different subpopulations of *M. lugubris*, we first ran F_1_-hybrid PSMC (hPSMC, [59]) using the same parameters as for the previous PSMC analyses and implementing 100 bootstraps replicates. Additionally, we estimated mitochondrial divergences within and between the two species and subpopulations of *M. lugubris* using the previously generated mitochondrial alignments. We applied the simple “2% rule” which assumes that in birds, the average divergence between two species is 2% per million years [60]. We compared these divergence time estimates with those obtained from the hPSMC analyses.

### Acoustic recordings and analysis

Acoustic recordings of 10 *M. gigantea* individuals and 28 *M. lugubris* individuals were obtained from an online repository of avian vocalizations (https://xeno-canto.org/), which covered different locations across New Guinea. We included all types of vocalisations-songs, calls and vocalisations of an unknown type in the analysis, unless the function of the vocalisation was specified by the recordist (eg: alarm). This is due to the high uncertainty in estimating the type of vocalisation in *M. lugubris,* and visual comparison between vocalisations classified as ‘songs’ versus ‘calls’ between individuals recorded in the same location, often showed that they were the same. The vocalisations of each individual (median = 9 vocalisations/individual) were measured by a single author (SR) using the Luscinia sound analysis program (version 2.17.11.22.01, [61]).

Each vocalisation was visualised using a Gaussian windowing function with the following spectrogram settings: 13 kHz maximum frequency, 5 ms frame length, 221 spectrograph points, 80% spectrograph overlap, 80 dB dynamic range, 30% dereverberation, and 50 ms of dereverberation range. Elements were measured as continuous sound traces and then grouped into syllables within each vocalisation (each vocalisation contained only one syllable).

The vocalisations were then compared using the dynamic time warping algorithm (DTW) in Luscinia, following the same settings used in Wheatcroft et al. [62] that has provided reliable grouping outputs for other songbird species. The final output of the DTW analysis was an acoustic dissimilarity matrix, from which we extracted 10 principal components using nonmetric multidimensional scaling.

## Results

Our evaluation of mapped reads against the *Corvus cornix* genome showed a median depth-of-coverage (DoC) of ∼ 9.252 *x* (min: 0.026 *x*, max: 30.935 *x*, SD: 7.444) and a median percentage of mapped reads at 89.5 % (min 0.2 %:, max: 95.8 %, SD: 23.693). Detailed values for each individual are listed in S1 Table.

During contamination control using mitochondrial assemblies, we observed an increased amount of heterozygote sites across the libraries in 6 individuals (S3 Table). Upon manual inspection using *Geneious Prime* we found that these heterozygote positions mostly appear in blocks and often within the same regions. This suggests that they were in fact nuclear mitochondrial sequences (NUMTs) that were wrongly mapped onto the mitochondrial genome instead of being a result of contamination. We also observed that non-reference alleles often appeared at a lower frequency (98.093 % of heterozygote sites had a reference allele frequency > 0.5, median reference allele frequency across all heterozygote sites at 0.874) and therefore disappeared during consensus calling, as the more frequent allele gets chosen during this step. Nonetheless, we manually excluded two regions from all samples with blocks (in total 5 700 bp out of the entire alignment’s 17 112 bp) of heterozygote sites shared across the majority of individuals. The remaining 11 412 bp were used to generate the mitochondrial phylogenies.

### Phylogenetic analyses on mitochondrial and nuclear DNA

We found high congruence between phylogenies built from mitochondrial and nuclear genomes (Fig. 1 A and S1 Fig.). Different window sizes and summary coalescent vs concatenated nuclear phylogenies also had little effect on the topology. We recovered three main clusters within *M. lugubris* that correspond to the geographic location of the samples on an east to west axis (Fig. 1). These clusters also align with previously described subspecies of *M. lugubris* [34]. The first cluster within *M. lugubris* consists of individuals inhabiting the Birds-Head of north-western New Guinea as well as an individual in the westernmost part of the Central Range. The next cluster inhabits the western and central parts of the Central Range of New Guinea, and the third cluster inhabits the eastern and south-eastern section of the Central Range as well as the isolated outlying Huon mountains. *M. gigantaea*, on the other hand, shows little differentiation between individuals compared to *M. lugubris*. Relationships within *M. gigantaea* are also in accordance with the geographical locality of the samples.

**Fig. 1.**
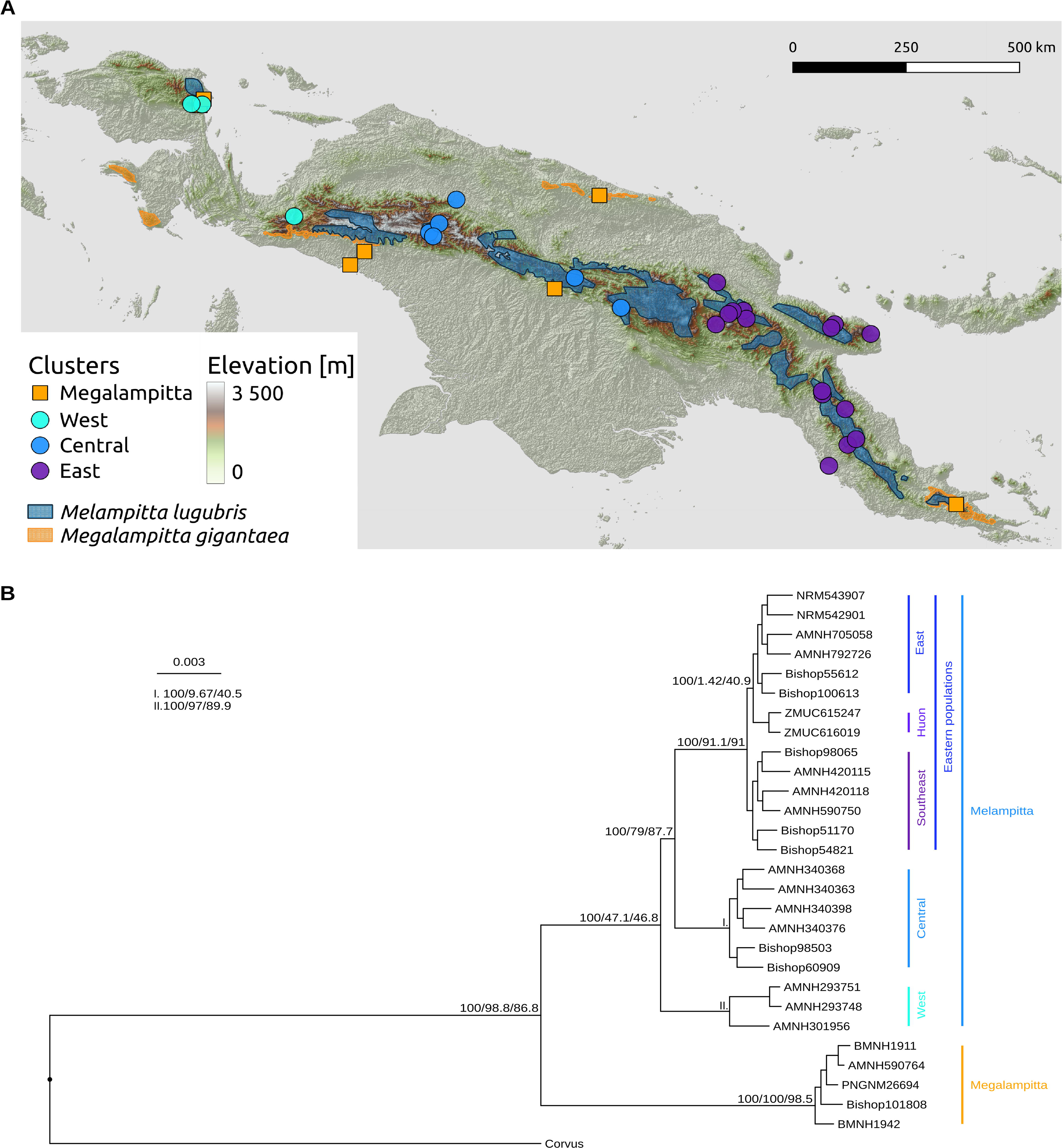
**A Distribution map and sampling sites** of Megalampitta gigantaea (orange distribution) and M. lugubris (blue distribution), subclusters of M. lugubris are also coloured differently. Shapefiles for administrative boundaries were obtained from geoBoundaries [63], the map was created using QGIS [64]. **B Nuclear phylogeny of Melampittidae** (based on concatenated 5 kbp window alignments) highlighting the subdivisions within M. lugubris (West, Central, East, Huon, Southeast), support values next to the main branches show Bootstraps/site concordance factors (sCF)/window concordance factors (wCF).

### Population structure and differentiation

We recover the same pattern of lower levels of differentiation in *M. gigantaea* compared to *M. lugubris* in the PCAs (Fig. 2 A), Heterozygosity (Fig. 2 B), nucleotide diversity (π, S4 Table, S2 Fig.) and admixture (Fig. 2 C). For *M. lugubris* Tajima’s D was consistently negative with a mean value of −0.902 (SD: 0.126, median: −0.897, S5 Table, S3 Fig.) which is indicative of either population expansion or a selective sweep. In *M. gigantaea* values for Tajima’s D were slightly above zero in the range of 0 – 0.2 (mean 0.103, SD: 0.040, median: 0.112, S5 Table, S3 Fig.). Positive values of Tajima’s D could indicate a reduction in population size or balancing selection acting, however as the values are so close to zero the population may just evolve neutrally. In the PCA (Fig. 2 A), PC1 separates the two species, afterwards *M. gigantaea* remains closely clustered up to PC4, while subgroups corresponding to geographic localities make up clusters within *M. lugubris* (S4 Fig.). The two distinct clusters of *M. lugubris* on PC2 (Fig. 2 A) separate eastern New Guinean populations and northwestern New Guinean populations as also observed in the phylogenetic tree (Fig. 1 B). Both heterozygosity and nucleotide diversity were significantly lower in *M. gigantaea* than in *M. lugubris*. Although we observed a clear trend of increasing heterozygosity with higher depth-of-coverage, the slopes for each population were similar and consistently higher in all but one population of *M. lugubris* (S5 Fig.). Admixture analysis revealed no substructure within *M. gigantaea* from K=2 to K = 7. For *M. lugubris* the observed clusters between K = 2-6 align with the clusters observed in the phylogenetic trees and in the PCAs. Further subdivisions within the main clusters of *M. lugubris* at higher values of K are also corresponding to the populations’ geographical location.

**Fig. 2.**
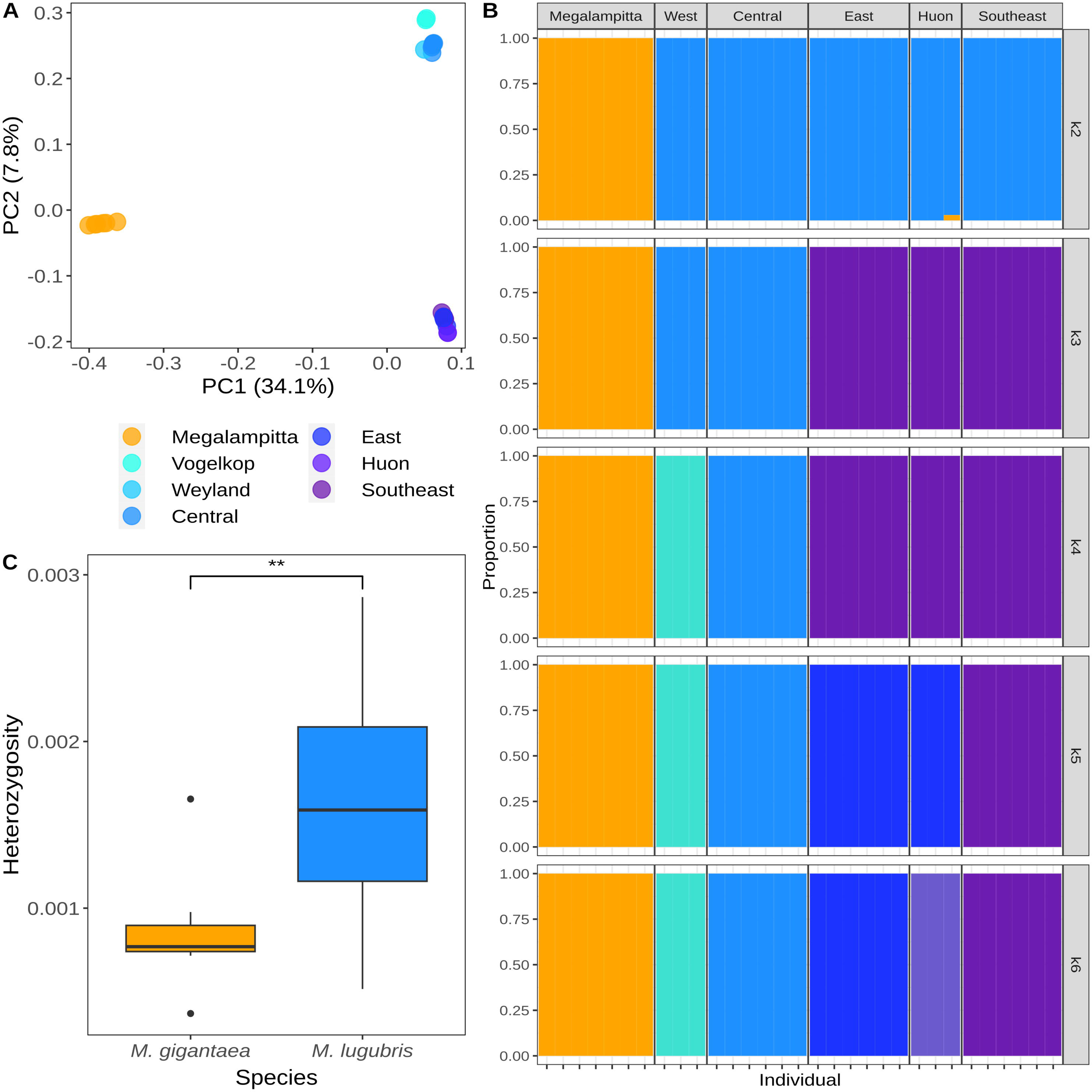
Genetic differentiation in Melampittidae. **A)** PCA showing the first two principal components for both species. Subclusters of Melampitta lugubris are also coloured differently. **B)** Admixture analysis from K = 1 to K = 6 **C)** Heterozygosity for all individuals between both species.

### Estimation of effective population size in time and divergence times

PSMC curves (Fig. 3) for samples from the same populations had similar shapes, but not entirely overlapping as depth-of-coverage varied between samples. Within *M. lugubris* the shape of the curves varied, but most of this variation could be ascribed to population specific events. In *M. gigantaea*, we observe an effective population size peak at around 200 Kya followed by a steady decline in effective population size up until around 40 Kya.

**Fig. 3.**
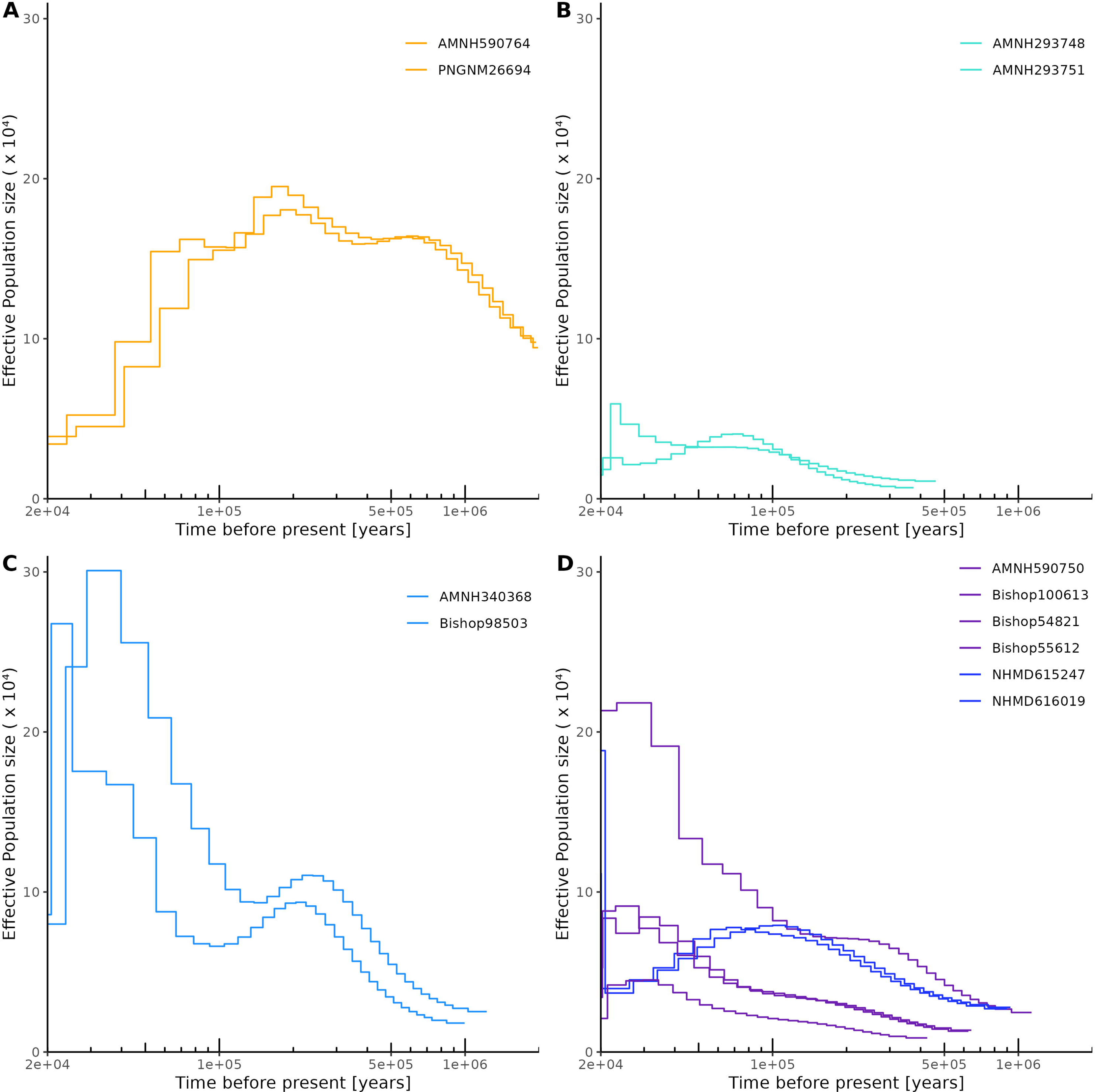
**PSMC plots** for two representative individuals of **A)** Megalampitta gigantaea **B)** Melampitta lugubris western population **C)** M. lugubris central population **D)** M. lugubris eastern population and Huon, colours in **D)** represent the Huon (blue) and (South)Eastern (purple) subclusters.

The divergence time obtained from hPSMCs curves for the split between *M. gigantaea* and *M. lugubris* was estimated to about 10 mya (S6 Fig.). Splits between subgroups within *M. lugubris* were estimated more recently with the split between Western+Vogelkop and Eastern populations at around 4-5 mya (S7 Fig.) and between Western and Vogelkop populations at about 3-4 mya (S8 Fig.). The next divisions within Eastern *M. lugubris* populations (East, Huon and Southeast) happened at similar times around 1 mya (S8 and S9 Figs.).

Mitochondrial divergences were, as expected, highest for comparisons between the *M. gigantaea* and *M. lugubris* and its subpopulations at a range of 9.8 – 13.3 %. Divergence within *M. gigantaea* was also lower (mean 0.912 %) than within *M. lugubris* (mean 4.979 %) or even in some of its subpopulations. For an extensive table with all comparisons of mitochondrial divergence see S11 and S12 Figs.. Divergence times obtained through the 2% rule were 4.9 – 6.6 mya for the split between *M. gigantaea* and *M. lugubris*, 3.7 – 5.6 mya for splits between Western+Vogelkop populations from Eastern populations of *M. lugubris* and Western populations from Vogelkop populations at 3.6 - 3.8 mya. Subdivisions within the Eastern populations were estimated at 0.2 – 2.7 mya.

### Acoustic recordings and analysis

The first ten principal components collectively explained 97% of the variation in vocalisations across the two *Melampitta* species. PC1, which explained 44.5% of the variation in all vocalisations, was more varied for *M. lugubris* (standard deviation (SD) = 0.086) compared to *M. gigantaea* (SD = 0.036). The same was true for PC2, where the standard deviation was once again higher for *M. lugubris* (SD = 0.11) compared to *M. gigantaea* (SD = 0.017). These results show that *M. lugubris* has greater acoustic diversity than *M. gigantaea* (Fig. 4).

**Fig. 4.**
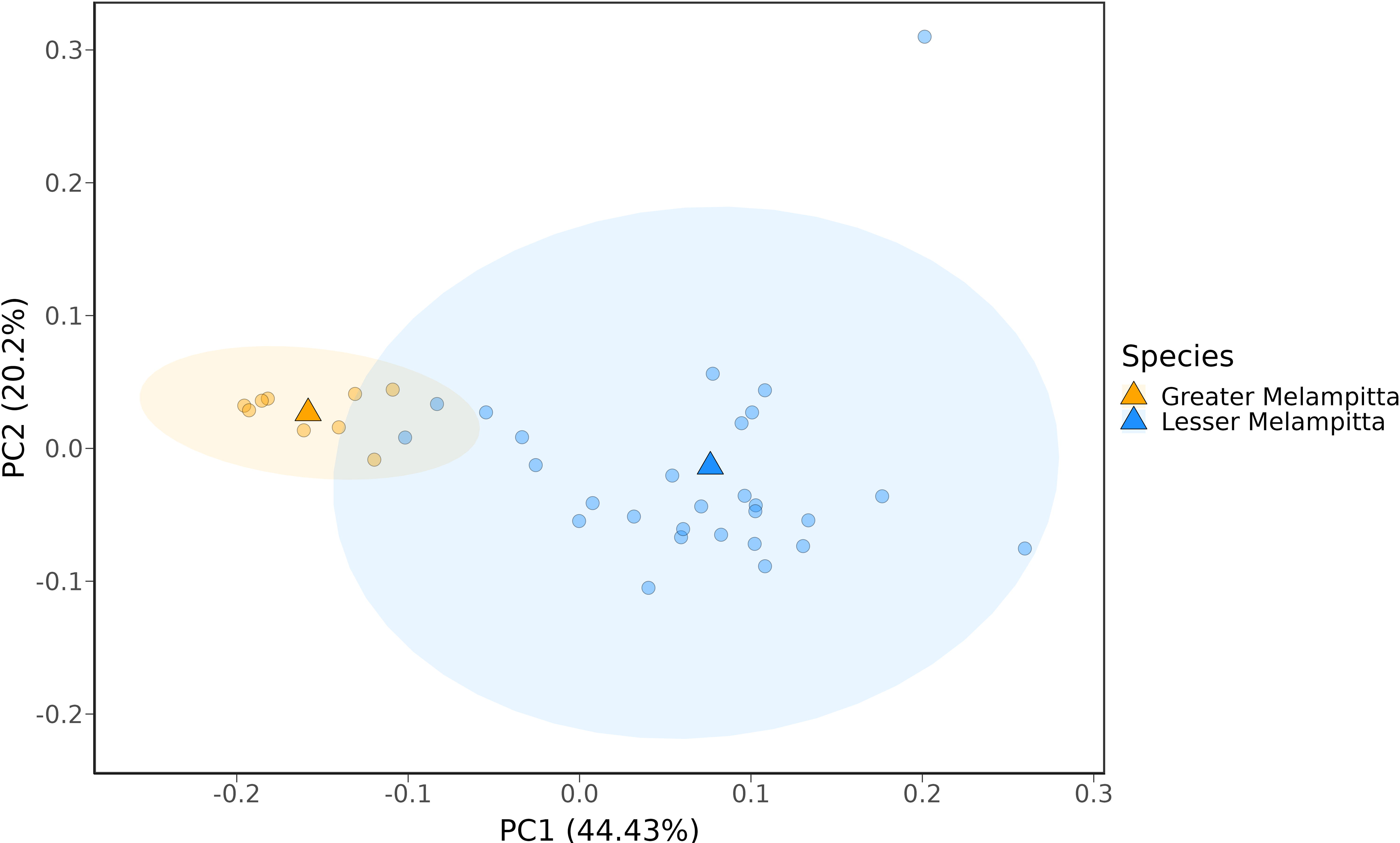
Acoustic variation across species: Principal component space (PC1-2) of vocalisations from M. gigantaea and M. lugubris. PC1 and PC2 scores are averaged within individuals and triangles represent species centroids. Ellipses contain 95% of vocalisations of each species. The Significant outlier within M. lugubris may represent an odd vocalisation that is not directly comparable with the other vocalisations included here. Note that vocalisations for M. gigantaea were only available from three localities (the Fakfak mountains in the Bird’s Neck, a locality in the southern Bird’s Neck and Tabubil in the central highlands) and vocalisations for M. lugubris were only available from two of the three distinct clades (samples were available from the western and central but no vocalisation data was available from the eastern and Huon populations).

## Discussion

The formation of the avifauna on New Guinea largely follows the predictions of taxon cycles [15,16] whereby new species form in or colonise through the lowlands and over time move upwards and become relictual at high elevations. The family Melampittidae is a species-poor old endemic lineage of New Guinea [31]. The family includes two extant species of which one (*Melampitta lugubris*) follows the general taxon-cycle expectation in that it is an old lineage that inhabits montane forests of New Guinea. The other species, *Megalampitta gigantea,* however, has a distribution associated with specific karst habitats at lower elevations and in foothills [34].

Our divergence time estimates suggest that *M. gigantea* and *M. lugubris* diverged from each other in the Miocene (at approximately 10 Mya based on hPSMC results), which is slightly younger than the divergence time estimated by Jønsson et al.[65] and slightly older than the divergence time estimated by McCullough et al. [66]. The three main populations of *M. lugubris* (Fig. 1 B) diverged from each other in the early Pliocene (at approximately 4-5 Mya based on hPSMC curves). A Pliocene divergence of *M. lugubris* populations coincides with major uplift of various mountain regions on New Guinea [67–69], which may have shaped the present population structure of *M. lugubris*. The distributional pattern of populations of *M. lugubris*, with one distinct Vogelkop population and a division of an eastern and a western population along the central mountain range, is also a pattern similar to that found in other New Guinean mountain birds with Pliocene divergences [30,70,71].

The PSMC curves of the three main populations of *M. lugubris* differ (Fig. 3), yet with a general trend of increasing population sizes towards the present. The exception to this is the population of the Huon mountains, which shows a continuous decrease in population size since approximately 100 Kya. Our interpretation is that eastern and south-eastern populations of the Central Range have maintained continuous gene flow, while the connectivity with the Huon population was broken or at least severely reduced as this population became isolated in the outlying Huon Mountain range.

Given a presumed poor dispersal capacity [33] and a patchy distribution at mid-elevations, we initially hypothesised that *M. gigantea* would exhibit a clear population structure. However, contrary to expectations, all our samples of *M. gigantea*, from localities scattered across New Guinea, cluster tightly together genetically (Fig. 1 and Fig**. *2***). Analysis of vocalisations also shows a similar pattern, as *M. gigantea* compared to *M. lugubris* exhibits less vocal differentiation (Fig. 4). This is fascinating and difficult to explain. Below, we discuss three scenarios that may provide possible explanations for these patterns. First, it is possible that continuous migration (or high rates of juvenile dispersal) of *M. gigantea* individuals maintains contact and gene flow between populations. However, an exclusively ground-dwelling lifestyle and the lack of long-distance flight capabilities, suggested by its morphology and field observations contradict this scenario [33,34]. Second, it is possible that their presently known fragmented distribution does not properly reflect their actual distribution, which may be more extensive [34,72]. Karst regions are generally species-poor in comparison to the species-rich tropical forests of New Guinea and such localised karstic areas dispersed throughout New Guinea may therefore have commanded less attention by ornithological surveys. Finally, it is possible that *M. gigantea* once had a wider more continuous distribution and that a recent decline has left scattered populations in small pockets of Karst habitat. The PSMC analyses support this scenario by showing that the population size of *M. gigantea* has dropped dramatically within the last 200 Ky (Fig. 3). The fact that *M. gigantea* is highly adapted to a very specific habitat type (nesting in deep holes in karst limestone that they have to climb out of [33] is, however, difficult to reconcile with this scenario. However, one may speculate that *M. gigantea* in the past had broader habitat preferences not only restricted to the present karst limestone habitats. Perhaps during the last 200 Ky, increased competition from other species forced *M. gigantea* to retract to a particular low-diversity habitat type, leaving behind the scattered distribution that we see today. Overall, we find it most plausible, that *M. gigantea* had a larger and more continuous distribution in the past, yet we acknowledge that the present distribution may be underestimated. Additional ornithological surveys to suitable habitats may, thus, reveal further *M. gigantea* populations.

## Conclusions

In this study, the rather surprising population structure of the two species of an old New Guinean avian family have been elucidated by genomic data largely obtained from historical museum collections. While the population structure of *Melampitta lugubris* is similar to those found in other mountain birds of New Guinea with similar age, the population structure of *Megalampitta gigantea* is intriguing. The study is an example of how intrinsic properties, such as those exhibited by *M. gigantea*, may cause their population dynamics to deviate from general biogeographical predictions. The study is also an example of how important museum collections are for increasing the knowledge of rare taxa that occur in remote regions. The levels of divergence between the three major populations of *M. lugubris* are well above those at which ornithologists would normally assign species rank. Consequently, we tentatively propose that these three populations should be elevated to species rank, *M. lugubris* (Schlegel, 1871) in the Vogelkop region, *Melampitta rostrata* (Ogilvie-Grant, 1913) in the western central range and *Melampitta longicauda* (Mayr & Gilliard, 1952) in the eastern central range.

## Acknowledgements

We thank all the staff and field assistants that facilitated fieldwork in Papua New Guinea. Notably, the Binatang Research Center and local communities from the villages of Towet and Yawan. We are also grateful for the assistance provided by the PNG National Museum and art gallery and the Conservation and Environment Protection Authority (CEPA) of Papua New Guinea for research permits and export permits. We would like to acknowledge the following museum collections and their managers that have generously provided us with tissue samples for this study: American Museum of Natural History, New York, USA (Paul Sweet, Pete Capainolo, Tom Trombone and Brian Smith); Bernice Pauahi Bishop Museum, Honolulu, USA (Molly Hagemann); Natural History Museum, London, UK (Robert Prys-Jones, Hein van Grouw and Mark Adams); Papua New Guinea National Museum and Art Gallery, Port Moresby, Papua New Guinea; Swedish Museum of Natural History, Stockholm, Sweden (Ulf Johansson) and the Natural History Museum of Denmark, Copenhagen, Denmark (Jan Bolding Kristensen). Computations were enabled by resources provided by the National Academic Infrastructure for Supercomputing in Sweden (NAISS) and the Swedish National Infrastructure for Computing (SNIC) at UPPMAX partially funded by the Swedish Research Council through grant agreements no. 2022-06725 and no. 2018-05973. Furthermore, the authors acknowledge support from the National Genomics Infrastructure in Stockholm funded by Science for Life Laboratory, the Knut and Alice Wallenberg Foundation and the Swedish Research Council, and SNIC/Uppsala Multidisciplinary Center for Advanced Computational Science for assistance with massively parallel sequencing and access to the UPPMAX computational infrastructure. Lastly, we would like to thank Nikolay Oskolkov for assistance with bioinformatic analyses as part of the drop-in services and the Swedish Bioinformatics Advisory Program of the National Bioinformatics Infrastructure Sweden (NBIS).

