## Supplementary material for "Species-specific dynamics may cause deviations from general biogeographical predictions – evidence from a population genomics study of a New Guinean endemic passerine bird family (Melampittidae)": S1 File

### PSMC background

The pairwise sequentially Markovian coalescent (PSMC) models variation in coalescent times to estimate changes in effective population size (1). Each diploid genome is a collection of hundreds of thousands of independent loci. By estimating the time to the most recent common ancestor (TMRCA) of the two alleles at each locus a distribution of TMRCA across the genome is created. Since the rate of coalescent events is inversely proportional to the effective population (N_e_) (2), PSMC identifies periods of change in the effective population size. For example, when many loci coalesce at the same time, it is a sign of a small N_e_ at that particular time. There are, however, considerable limitations of the PSMC method. For example, PSMC cannot recover recent changes in N_e_, i.e. younger than 10,000 years as well as sudden changes in N_e_ as it may smooth out curves depending on chosen interval parameters (1). This method is based on several assumptions such as an absence of selection or random mating that may not be applicable to natural populations and will therefore lead to false estimations of N_e_ (3,4). Also, it is sensitive to estimates of the generation time and average mutation rate as these are used to scale the TMRCA distribution into years. However, the generation time and mutation rate estimates do not change the shape of the PSMC curve but rather push the PSMC curves along the axes. For example, a halved generation time will double the estimate of N_e_ (given a fixed mutation rate per year), and a halved mutation rate per year will move the curve backward in time as well as double the estimate of N_e_. While we, therefore, cannot rely on precise estimations of effective population sizes at exact time points, we can still use the estimates in a broader and relative comparison.
