## Supplementary material for "Species-specific dynamics may cause deviations from general biogeographical predictions – evidence from a population genomics study of a New Guinean endemic passerine bird family (Melampittidae)": S2 File

### Codes and parameters

**Mitochondrial alignment:**

The alignment was generated using mafft *v7.407* [1] with the following parameters:

mafft --thread 16 --reorder --adjustdirection –globalpair

--maxiterate 1000 $IN

**Parameter explanation:**

- -thread 16: Number of threads used
- -reorder: Order the output based on alignment similarity
- -adjustdirection: Generate reverse complements and align them
- –globalpair: Apply Needleman-Wunsch algorithm to generate a global alignment
- -maxiterate 1000: Maximum number of iterations. Recommended when

using -globalpair

**ANGSD**

Beagle files were generated using ANGSD *v0.938* [2] using these parameters:

angsd -bam Bams.list -out $OutPath -doSaf 1 -GL 1 -doGlf 2 -doMajorMinor 1 -ref $REF -anc $REF -doMaf 1 -minMaf 0.05 -SNP_pval 1e-6 -doCounts 1 -setMinDepth 62 -setMaxDepth 465 -minInd 15 -minQ 20 -minMapQ 20 -uniqueOnly 1 -only_proper_pairs 1 -remove_bads 1 -baq 1 -C 50 -P 10

**Parameter explanation:**

- -bam bamlist.txt: Specify we're using bam, give the name of the bamlist
- -out: set prefix for output files
- -doSaf 1: Calculate the Site allele frequency likelihood based on individual genotype likelihoods assuming Hardy-Weinberg-Equilibrium
- -GL 1: Estimate genotype likelihoods using SAMtools model
- -doGlf 2: Generate beagle input file
- -doMajorMinor 1: Infer major and minor from genotype likelihoods
- -ref $REF: Give reference sequence
- -anc $REF: Give ancestral sequence, as we did not have one available, we used the reference sequence, but following steps need to be performed with -fold 1
- -doMaf 1: Estimate allele frequencies with known major minor
- -minMaf 0.05: Only work with sites with a minor allelic frequency above 0.05
- -SNP_pval 1e-6: Test for polymorphic sites and only output the ones which have a likelhood ratio test p-value < 1e-6
- -doCounts: Output the counts of the different bases

-setMinDepth 62: Discard sites if their total sequencing depth (all individuals added together) is below the given value. As an example I chose 62 (2*31) to get on average at least 2 reads per site per individual for this dataset (n = 31 individuals)

-setMaxDepth 465: Discard sites if their total sequencing depth (all individuals added together) is above the given value. E.g. for the same subset 465 (31*15), the factor is an arbitrary decision, which should removes repetitive sites that can cause excessive depth

-minInd 15: Only keep sites with at least minIndDepth (default is 1) from at least the given number of individuals, I chose 15 to include at least half of the individuals

-minQ 20: Minimum allowed base quality score

-minMapQ 20: Minimum allowed mapping quality score

-uniqueOnly 1: Remove reads that have multiple best hits.

-only_proper_pairs 1: Include only proper pairs (pairs of read with both mates mapped correctly)

-remove_bads 1: Same as samtools’ -x flag which removes reads with a flag above 255 (not primary, failure and duplicate reads)

-baq 1: Perform base alignment quality (BAQ) computation. Reduces false SNP calling at possibly misaligned bases [3]

-C 50: Adjust mapping quality among reads with high numbers of mismatches. For reads mapped with BWA, a value of 50 is recommended (according to samtools’ documentation)

-P 10: Number of threads to be used

**PCA**

Covariance matrices for PCAs were generated using PCANGSD [4]

pcangsd.py -beagle ${InPATH}/${i}.beagle.gz -threads 8 -o ${OutPATH}/${i}

**Parameter explanation:**

- -beagle: Specify input beagle file
- -threads 8: Number of cores to be used
- -o: Specify output directory and name

**NGSAdmix**

Admixture proportions were estimated through NGSAdmix [5] with the following script:

```
#Loop through values of k
for k in $(seq 1 $MaxK)
do
  #Loop through number of replicates (also used as seed)
  for n in $(seq 1 $MaxReps)
  do
    angsd/misc/NGSadmix -likes ${InPATH}/${i}.beagle.gz -seed ${n} -K ${k} -P 4 -o ${OutPATH}/${i}/${i}_k${k}_r${n}
  done
done
```

**NGSadmix parameter explanation:**

-likes: Input file in .beagle format

-seed ${n}: Initial seed to be used in the expectation-maximisation (EM) algorithm. The number of replicate (e.g 1-10) is being used as seed.

-K ${k}: Number of ancestral populations

-P 4: Number of threads

-o: Output path and prefix

**Heterozygosity estimates**

To obtain estimates of individual heterozygosity, we first generated sample allele frequency (saf) files using ANGSD with the following settings:

angsd -i $bam -out $OutPath/${filename} -doSaf 1 -GL 1 -doMajorMinor 1 -ref $REF -anc $REF -doMaf 1 -doCounts 1 -setMinDepth 2 -setMaxDepth 15 -minQ 20 -minMapQ 20 -uniqueOnly 1 -only_proper_pairs 1

-remove_bads 1 -baq 1 -C 50 -P 10

**Parameter explanation (missing parameters are explained above under ANGSD):**

-i: Specify single input .bam file

In the next step, .saf files were converted into global site frequency spectra (sfs):

angsd/misc/realSFS ${i} -P 16 -fold 1 > ${OutPATH}/${filename}.sfs

**Parameter explanation:**

- -fold 1: Generate a folded site frequency spectrum. Necessary when no ancestral sequence is available and the reference is used in its stead
- -P 16: Number of threads

**Thetas estimation:**

SAFs were generated the same way as for NGSAdmix, but filters for p-value or MAF were left out, the applied parameters were as follows (see above for explanations):

angsd -bam Bams.list -out $OutPath -doSaf 1 -GL 1 -doGlf 2 -doMajorMinor 1 -ref $REF -anc $REF -doMaf 1 -doCounts 1 -setMinDepth 62 -setMaxDepth 465 -minInd 15 -minQ 20 -minMapQ 20 -uniqueOnly 1 -only_proper_pairs 1 -remove_bads 1 -baq 1 -C 50 -P 10

SFS were generated with the same command as for single individuals:

angsd/misc/realSFS ${InPATH}/${i}.saf.idx -fold 1 -P 12 > ${OutPATH}/${i}.sfs

Chromosome-wide theta estimates were obtained using the following commands:

angsd/misc/realSFS saf2theta ${i}.saf.idx -sfs ${i}.sfs -fold 1 \

-P 1 -outname ${OutPATH}/${i}

angsd/misc/thetaStat do_stat ${OutPATH}/${i}.thetas.idx

**Parameter explanation:**

- -fold 1: Generate a folded site frequency spectrum. Necessary when no ancestral sequence is available and the reference is used in its stead
- -P 1: Number of threads
- -outname: Specify output path and prefix

Additionally, we obtained window-based thetas with the following command:

angsd/misc/thetaStat do_stat ${OutPATH}/${i}.thetas.idx -win 10000 -step 20000 -outnames ${OutPATH}/${i}.Win

**Parameter explanation:**

- -win 10000: Define sliding window size
- -step 20000: Define step size in which windows are moved
- -outnames: Specify output path and filename
