## Supplementary figures and images for "Species-specific dynamics may cause deviations from general biogeographical predictions – evidence from a population genomics study of a New Guinean endemic passerine bird family (Melampittidae)"

### S1.tiff

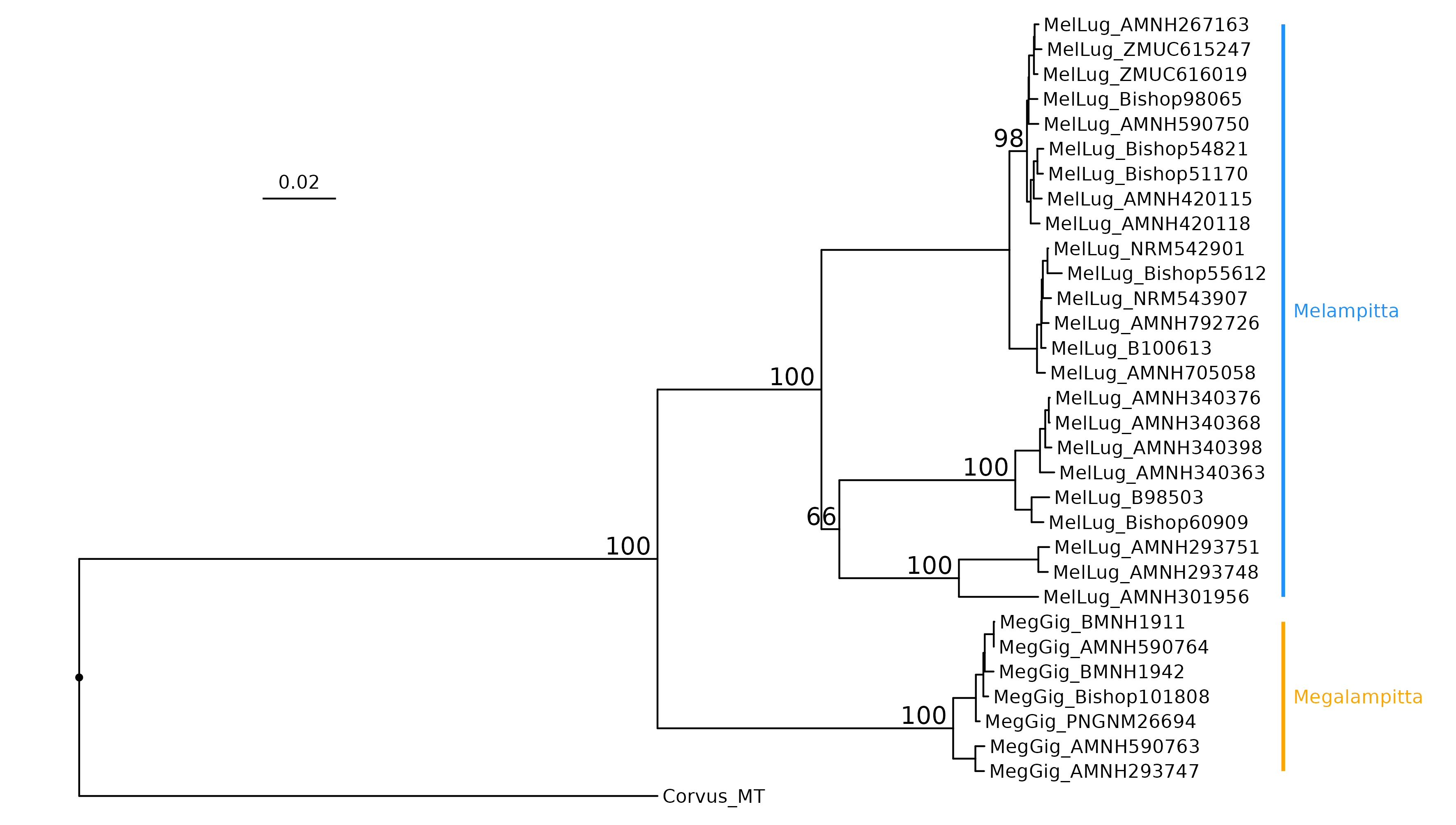

### S2.tiff

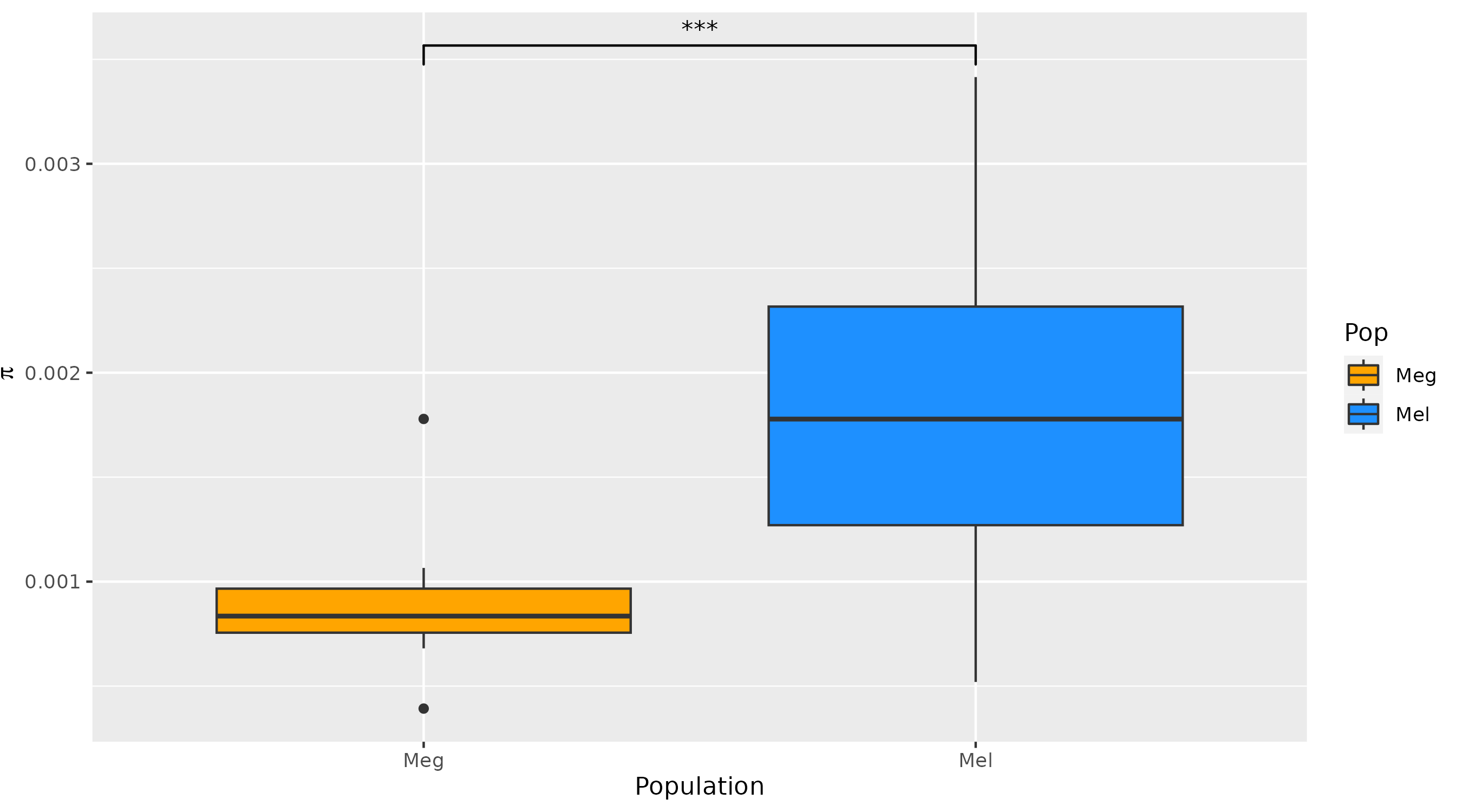

### S3.tiff

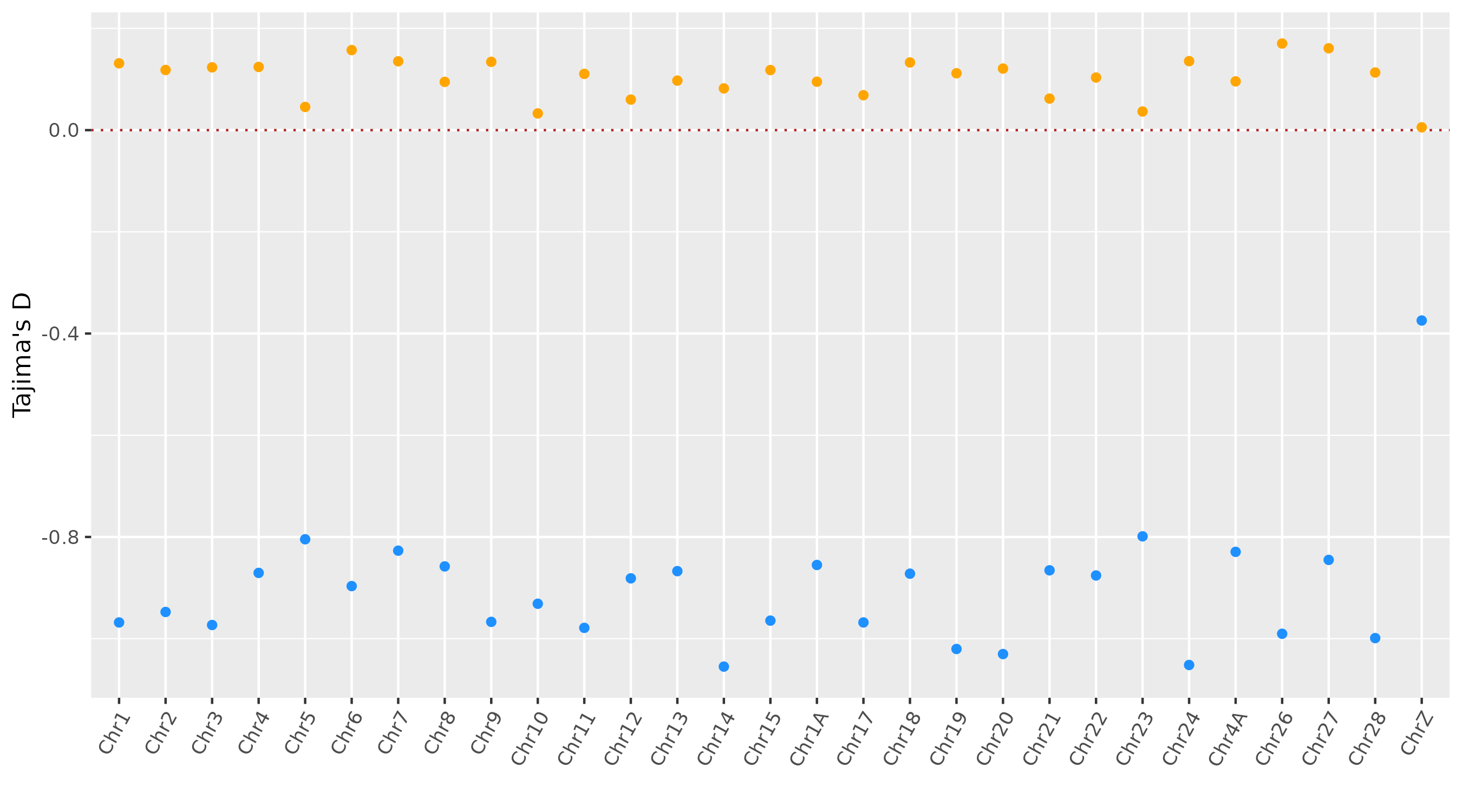

### S4.tiff

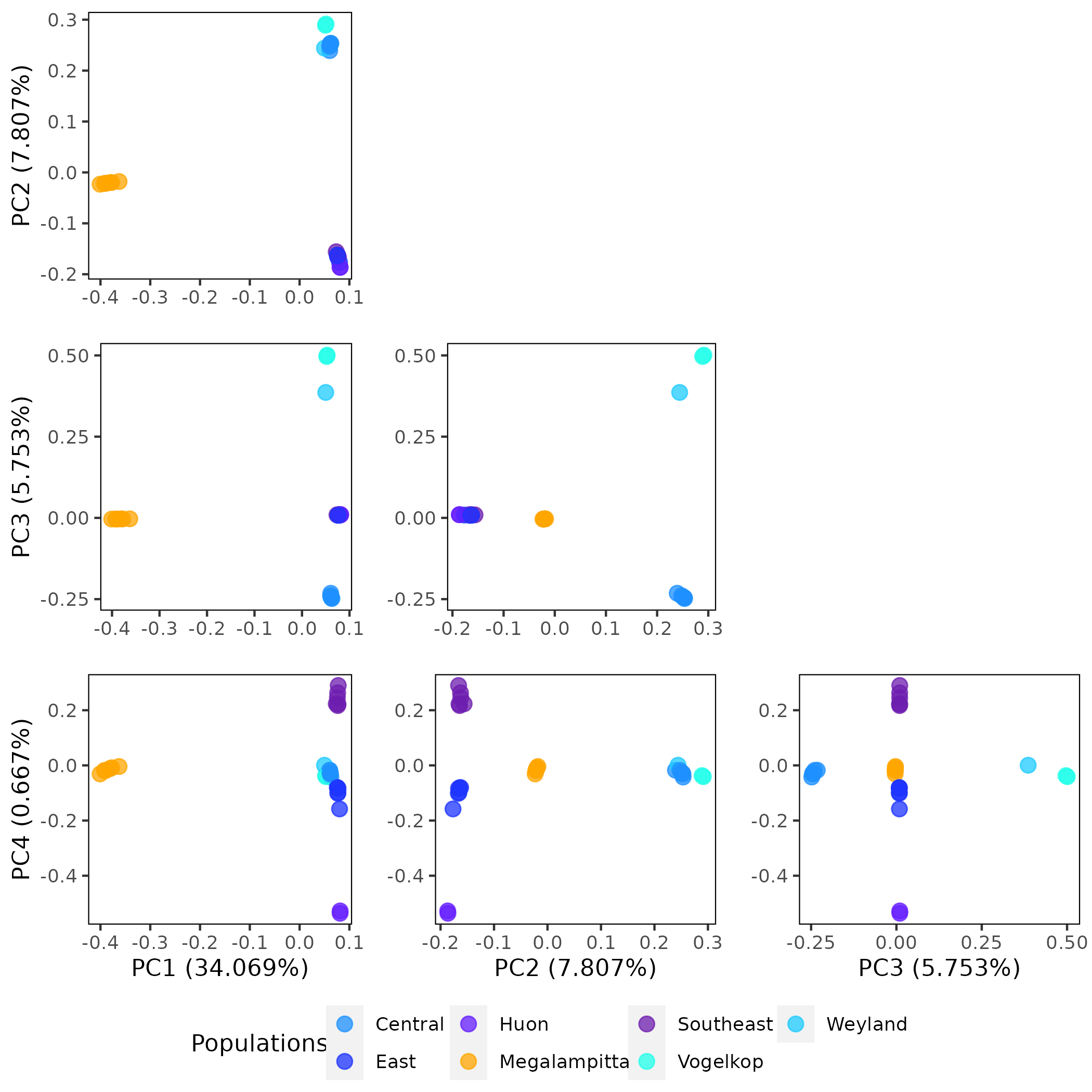

### S5.tiff

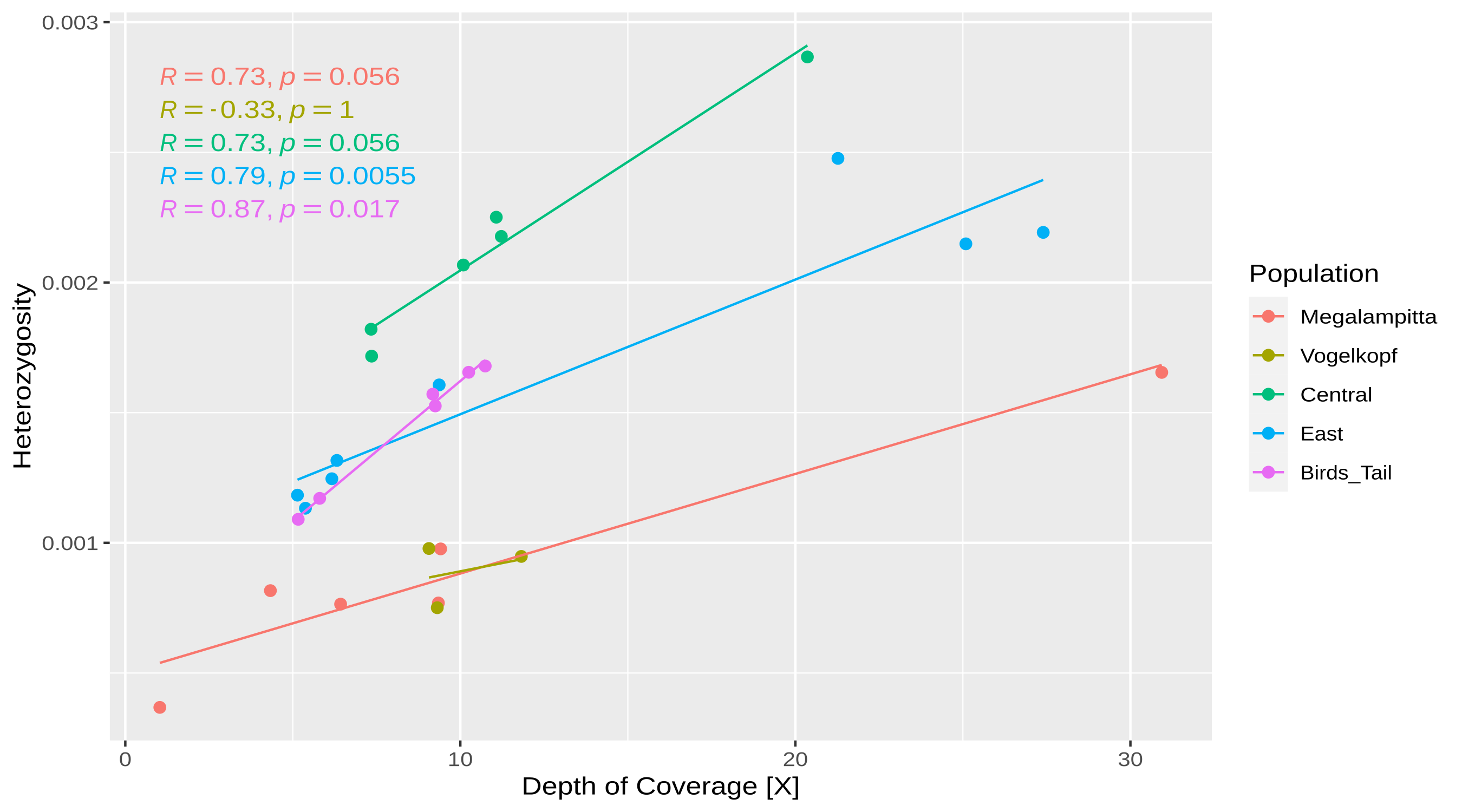

### S6.tiff

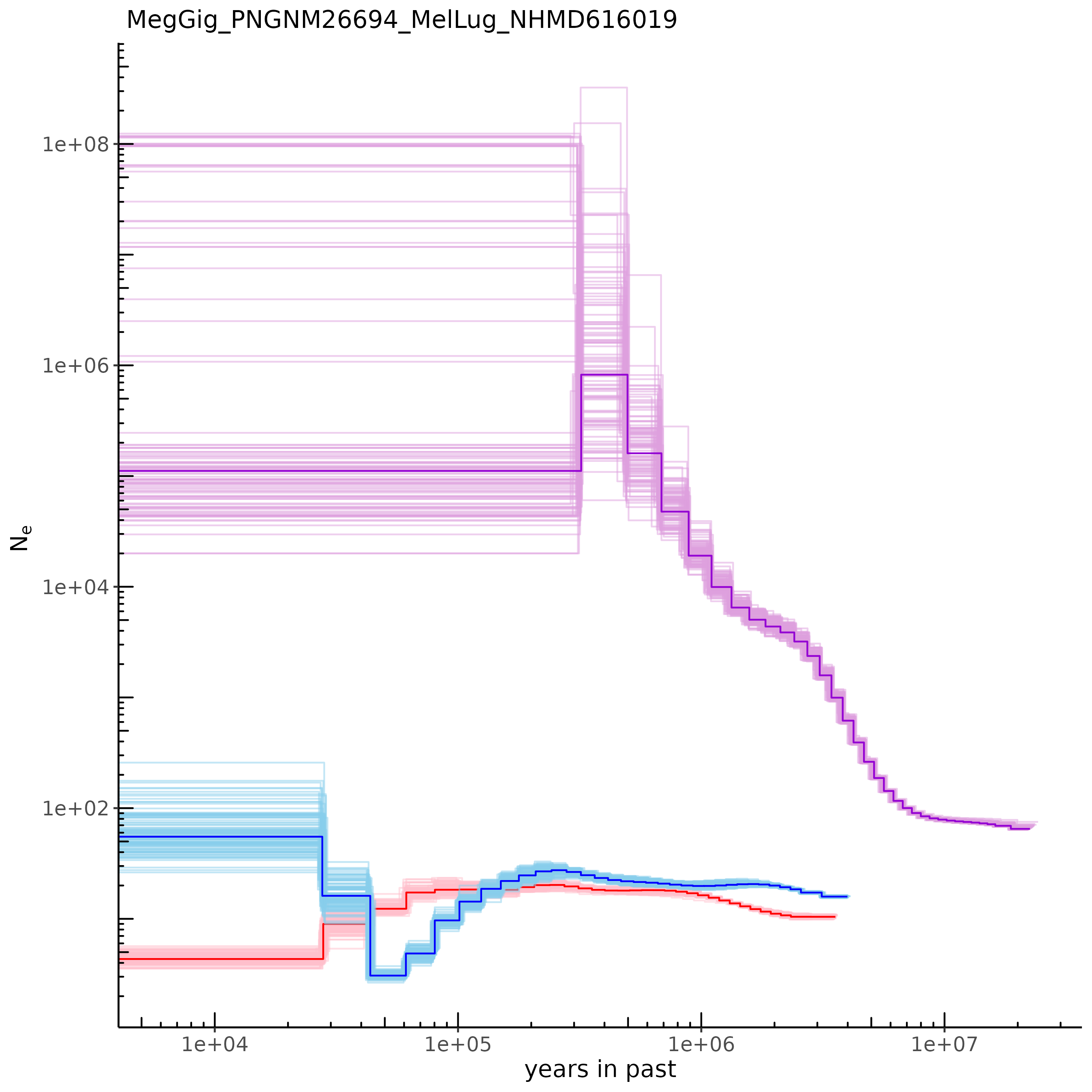

### S7.tiff

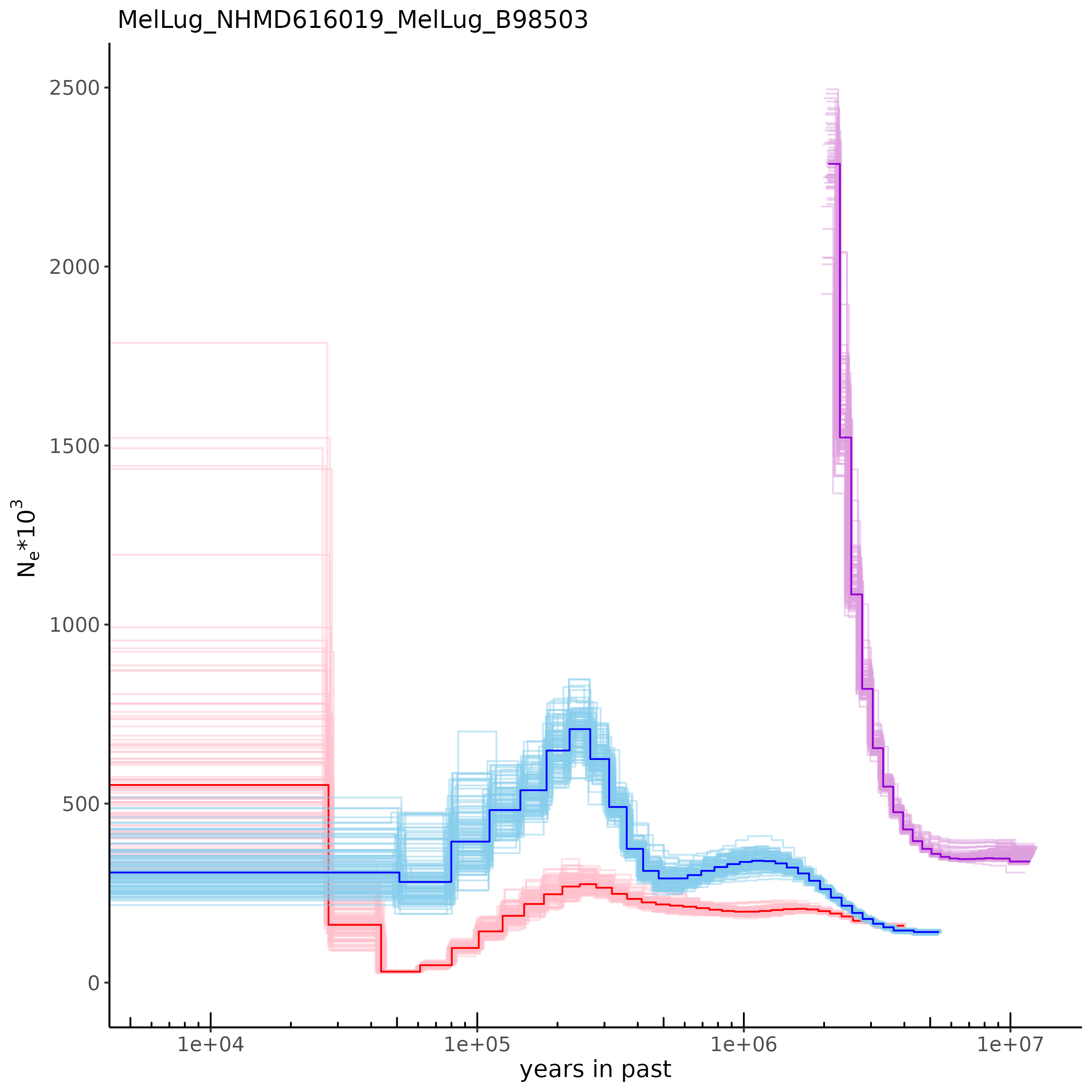

### S8.tiff

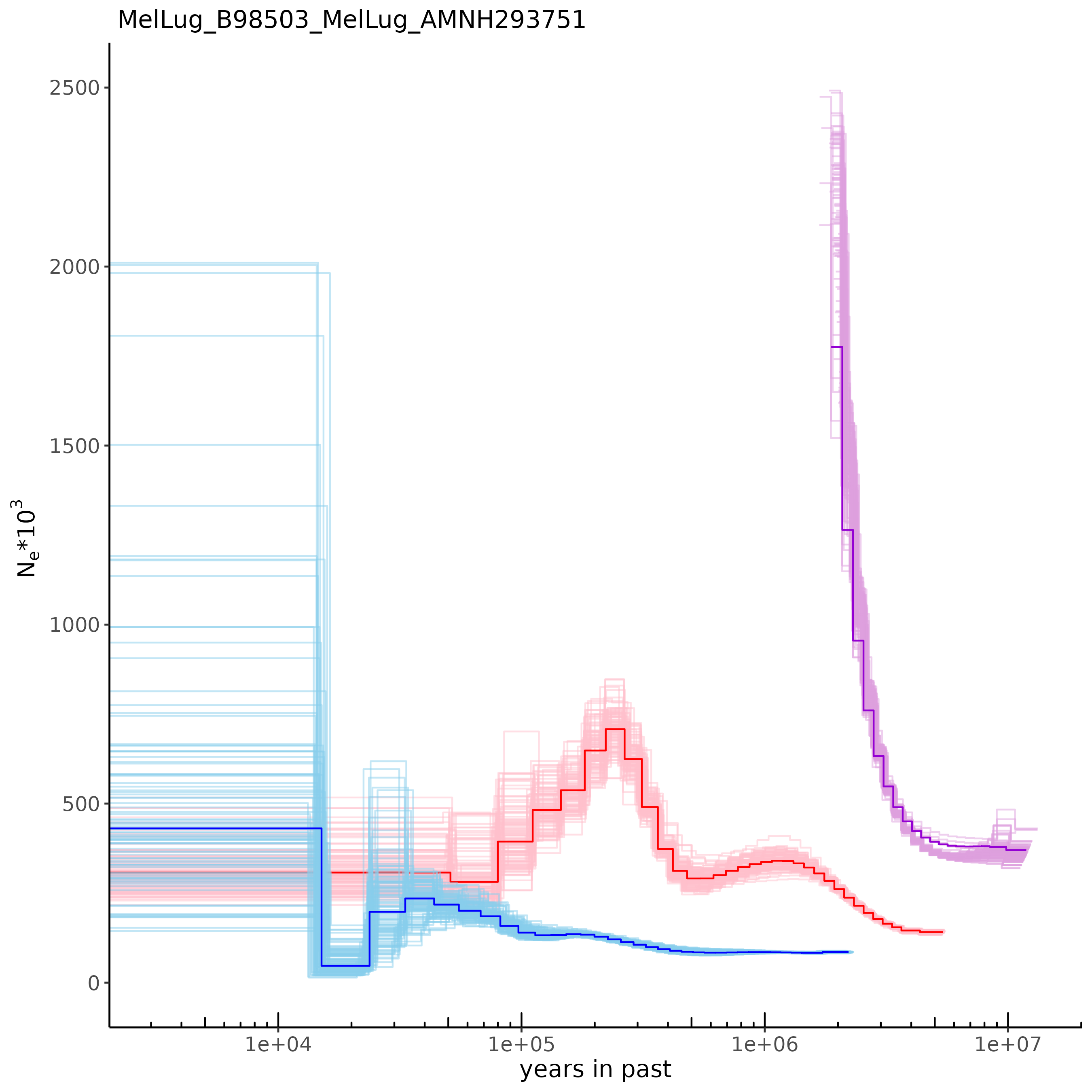

### S9.tiff

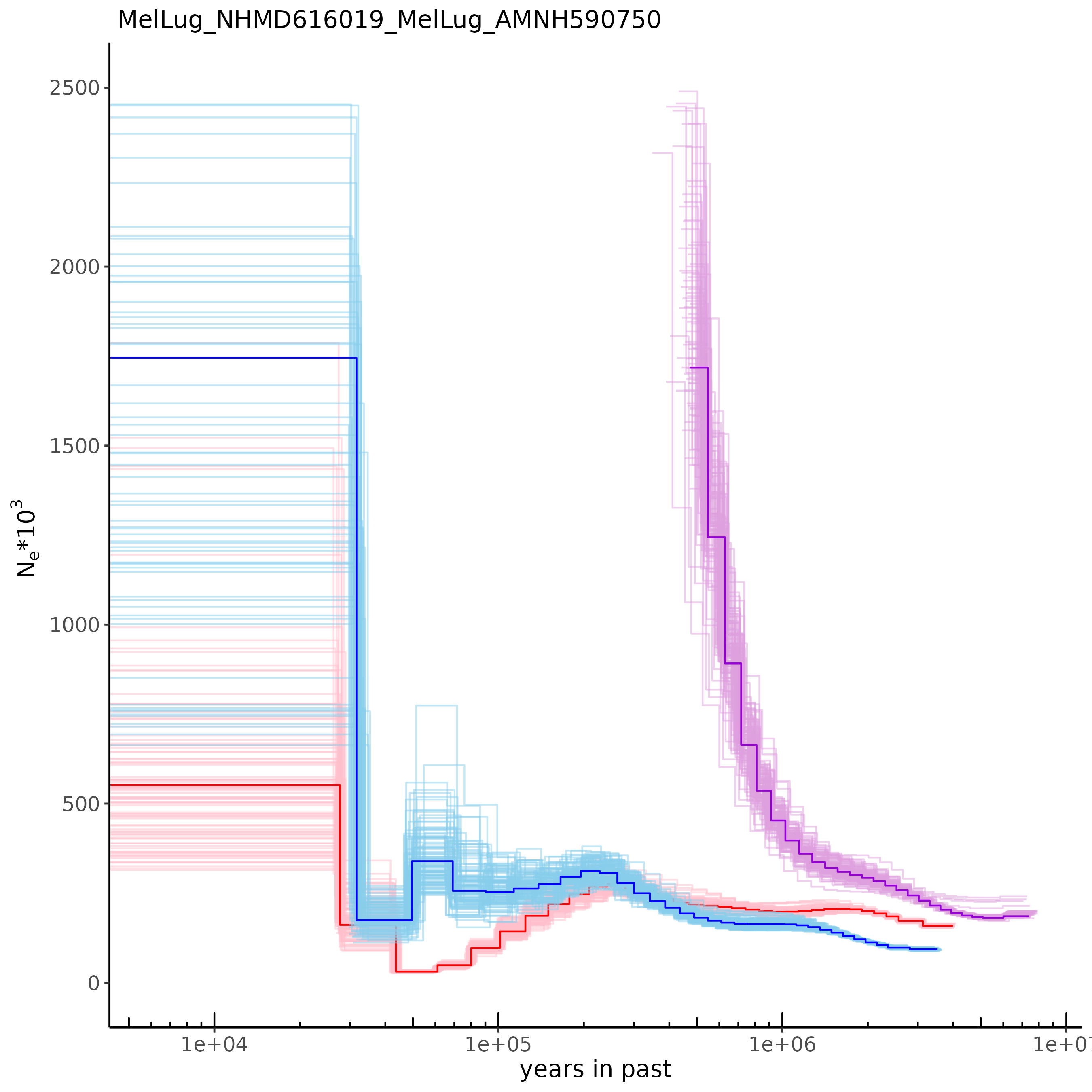

### S10.tiff

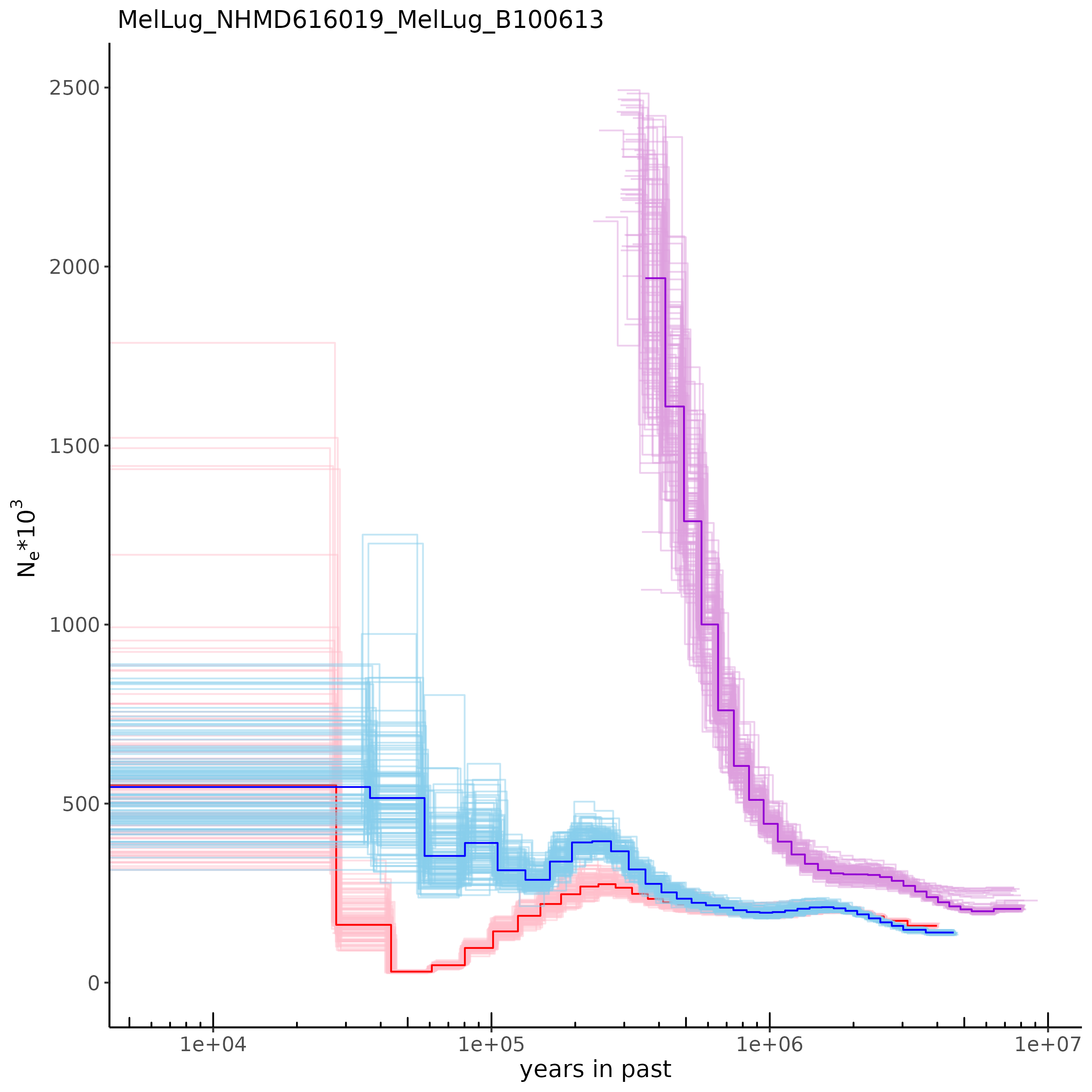

### S11.tiff

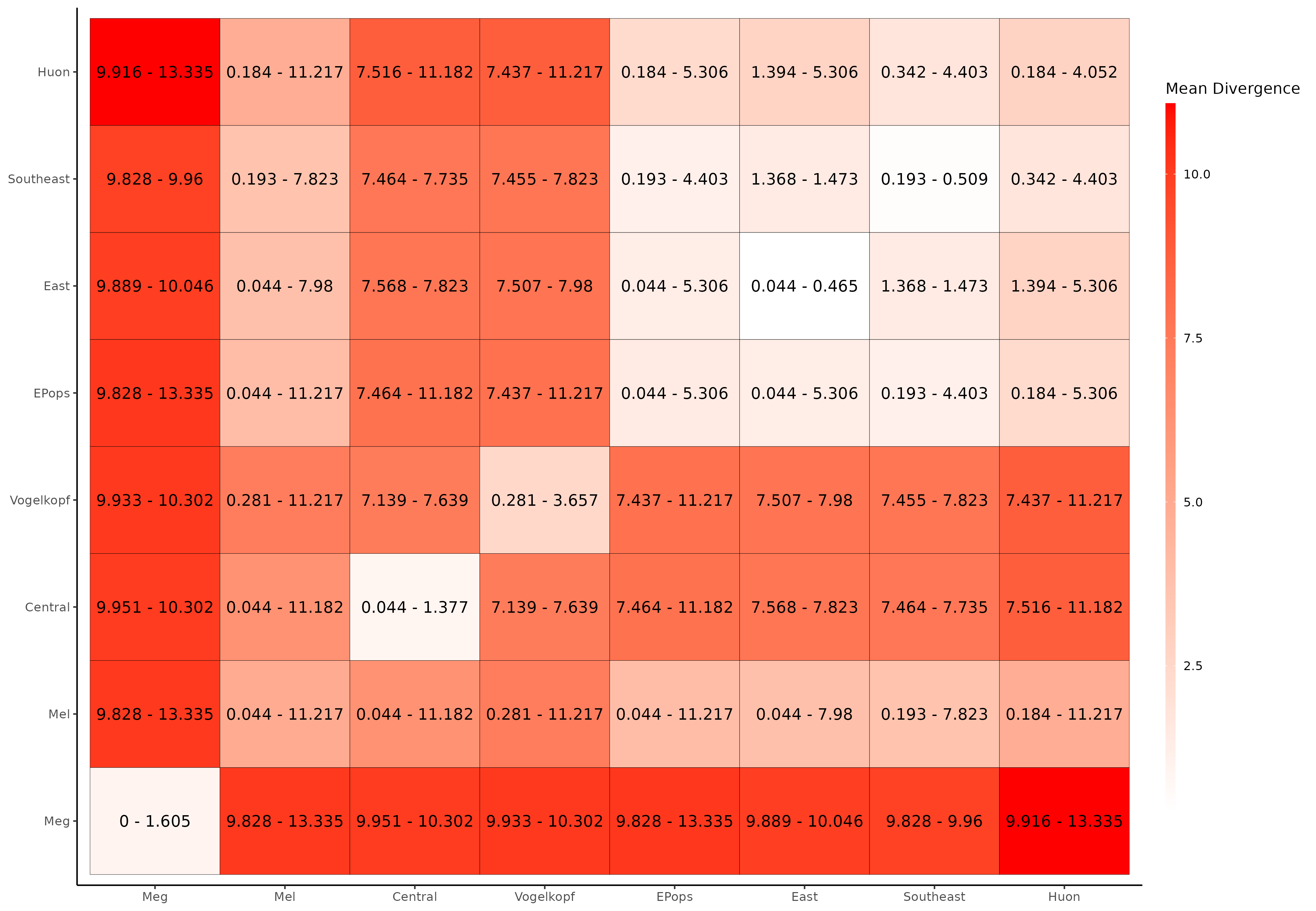

### S12.tiff

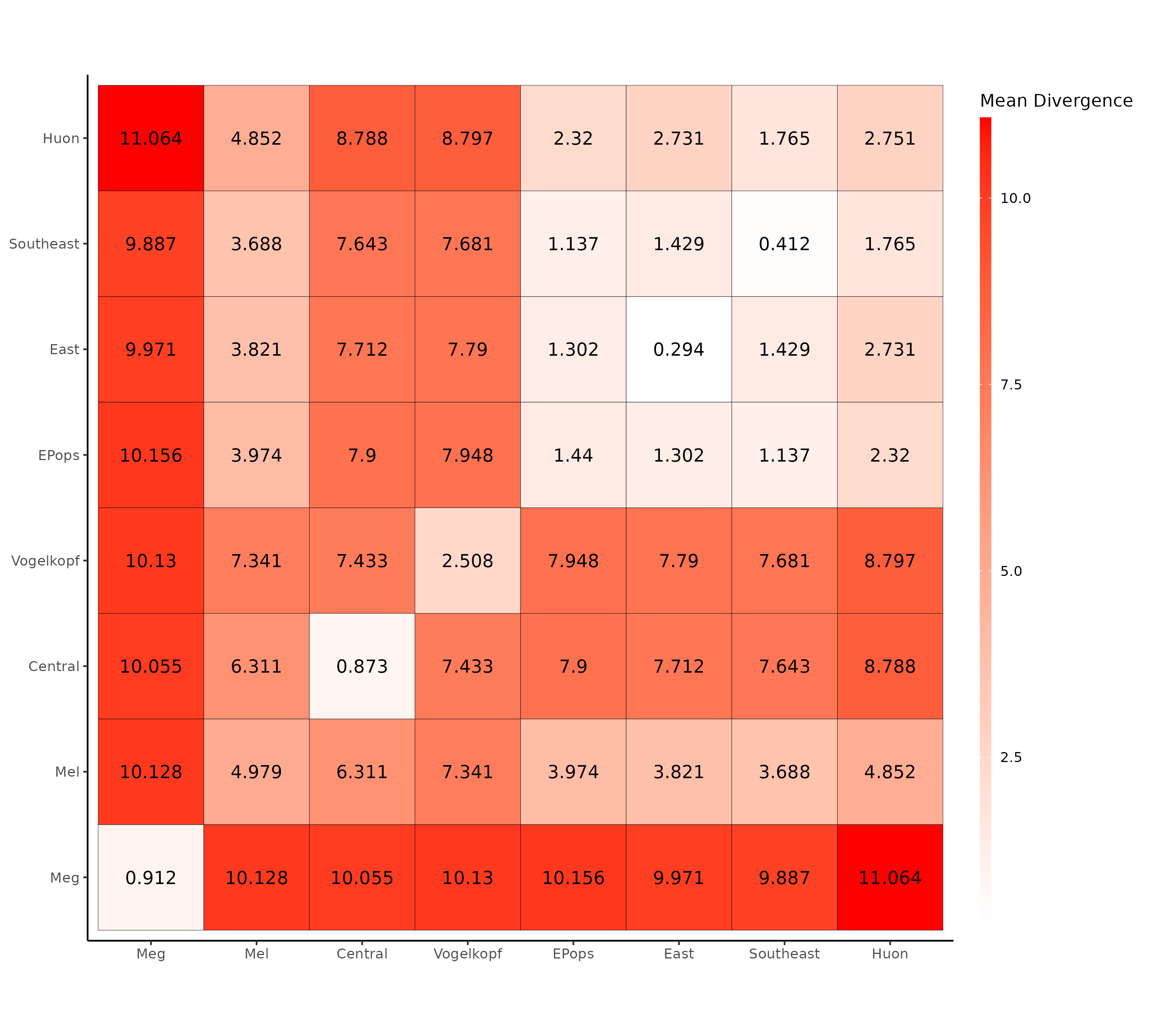
